# ISGylation of the SARS-CoV-2 N protein by HERC5 impedes N oligomerization and thereby viral RNA synthesis

**DOI:** 10.1101/2024.05.15.594393

**Authors:** Junji Zhu, Guan Qun Liu, Christopher M. Goins, Shaun R. Stauffer, Michaela U. Gack

## Abstract

Interferon (IFN)-stimulated gene 15 (ISG15), a ubiquitin-like protein, is covalently conjugated to host (immune) proteins such as MDA5 and IRF3 in a process called ISGylation, thereby limiting the replication of Severe acute respiratory syndrome coronavirus 2 (SARS-CoV-2). However, whether SARS-CoV-2 proteins can be directly targeted for ISGylation remains elusive. In this study, we identified the nucleocapsid (N) protein of SARS-CoV-2 as a major substrate of ISGylation catalyzed by the host E3 ligase HERC5; however, N ISGylation is readily removed through de-ISGylation by the papain-like protease (PLpro) activity of NSP3. Mass spectrometry analysis identified that the N protein undergoes ISGylation at four lysine residues (K266, K355, K387 and K388), and mutational analysis of these sites in the context of a SARS-CoV-2 replicon (N-4KR) abolished N ISGylation and alleviated ISGylation-mediated inhibition of viral RNA synthesis. Furthermore, our results indicated that HERC5 targets preferentially phosphorylated N protein for ISGylation to regulate its oligomeric assembly. These findings reveal a novel mechanism by which the host ISGylation machinery directly targets SARS-CoV-2 proteins to restrict viral replication and illuminate how an intricate interplay of host (HERC5) and viral (PLpro) enzymes coordinates viral protein ISGylation and thereby regulates virus replication.

## Introduction

Severe acute respiratory syndrome coronavirus 2 (SARS-CoV-2), a beta-coronavirus that emerged in late 2019 and continues to spread across the globe, causes mild to severe symptoms and clinical manifestations known as coronavirus disease 2019 (COVID-19) (1–3). SARS-CoV-2 encodes for ∼30 proteins in its +ssRNA genome, and virus replication and transcription are exquisitely orchestrated by a number of viral and host proteins (2, 4). RNA transcription and replication takes place at the viral replication and transcription complex (RTC), which is localized to unique double-membrane vesicles (DMVs) inside infected cells (2, 5, 6). Among the SARS-CoV-2 proteins crucial for viral transcription and replication, the nucleocapsid (N) protein stands out as a pivotal player. This 419-amino-acid long protein exhibits RNA-binding and dimerization capabilities essential for various stages of the viral life cycle (7, 8). Upon SARS-CoV-2 entry into the host cell, the N protein is dissociated from the viral RNA genome, initiating the viral gene expression program. This process is tightly regulated in a spatial and temporal manner to ensure efficient replication. Specifically, the N protein orchestrates the RTC where it regulates transcription and replication processes (2). At later stages of infection, the N protein is essential for packaging viral RNA for export into newly formed virions (9). Apart from its fundamental roles in the viral life cycle, the N protein has been proposed to act as a viral inhibitor of RNA interference (RNAi), a host defense mechanism that targets viral RNA for degradation, thus promoting viral replication (10). Furthermore, SARS-CoV-2 N has been reported to manipulate multiple immune signaling pathways and proteins (such as Gasdermin D, NLRP3, RIG-I, MAVS and IFNAR signaling) (11–16). Accumulating evidence suggests that the multi-faceted functions of coronaviral N require tight regulation by several posttranslational modifications (PTMs) including multi-site phosphorylation, methylation and ubiquitination (among other PTMs) (17–25); however, how ubiquitin-like proteins regulate SARS-CoV-2 N activity is elusive.

IFN-stimulated gene 15 (ISG15) is unique among the many ISGs because it functions as a posttranslational modifier conjugated to a myriad of proteins, thus determining the fate of many host cellular pathways (26, 27). At its basal expression levels or following transcriptional upregulation in response to type I or III IFN stimulation (i.e., IFN-β as well as IFN-α and IFN-λ subtypes), ISG15 can be conjugated to specific lysine residues in substrate proteins, an activity mediated by an E1 enzyme (UBA7/Ube1L), an E2 conjugating enzyme (Ube2L6/UbcH8), and certain ligase enzymes or E3s (such as, HERC5, ARIH1 and EFP/TRIM25) that conjugate ISG15 to the substrate (28–33). Inversely, deISGylases – of note, both host (e.g., USP18, also called UBP43) and viral (e.g., papain-like proteases of *Coronaviridae* family members) deISGylating enzymes exist – reverse ISGylation events and thereby counteract the regulatory role of ISGylation on the substrate protein (34–36). In addition to being covalently conjugated to target proteins, which is mediated by a diglycine motif (LRGG) in ISG15, ISG15 can also bind to proteins noncovalently and thereby modulate the activity of the interacting protein (37, 38).

ISG15 has been shown to mediate both antiviral and proviral roles (39–41). Covalent ISGylation of specific receptors or signaling mediators in innate sensing pathways promotes antiviral signaling (41). For instance, the cytoplasmic-localized dsRNA receptor MDA5 is covalently modified with ISG15 in its CARD domains, which helps the sensor protein to form higher-order assemblies and thereby promotes antiviral gene induction to restrict a range of RNA viruses including SARS-CoV-2 (35). Similarly, cGAS and its adaptor protein STING, as well as the downstream transcription factor IRF3, undergo ISGylation in infected cells, which potentiates antiviral signaling (42–45). On the other hand, ISG15 in its unconjugated form exerts a proviral activity by dampening signaling by the type I IFN receptor (IFNAR), an effect underlying certain in-born ISG15 deficiencies in humans (40). Recent studies indicated that ISG15 can be secreted from infected cells and instigates immunomodulatory events akin to a cytokine or extracellular signaling mediator (46). Beyond innate immunity, ISGylation also regulates the activity, stability, subcellular localization or interactome of other host-cell proteins, thereby playing a pivotal role in fundamental processes such as DNA damage, autophagy or translation (27, 47).

Not surprisingly, viral proteins once produced in the infected cell can also be conjugated with ISG15 by the host E1-E2-E3 ISGylation machinery. However, as compared to our knowledge about ISG15’s role in regulating host cellular processes, relatively little is known about how ISGylation modulates the activities of viral proteins, in particular coronaviral proteins and thereby the coronavirus lifecycle.

Here, we show that the oligomeric state of the SARS-CoV-2 N protein is regulated by ISGylation mediated by cellular HERC5, which leads to restriction of viral RNA synthesis. We further identified an intricate crosstalk between N ISGylation and phosphorylation regulating N oligomerization and thereby its function.

## Results

### The SARS-CoV-2 N protein undergoes ISGylation, which is regulated by PLpro’s de-ISGylating activity

ISGylation, the covalent conjugation of ISG15, regulates a myriad of cellular processes including innate immunity to SARS-CoV-2 (and other virus) infections (35, 45); however, the extent to which proteins encoded by SARS-CoV-2 are direct targets of ISGylation remains still unknown. To address this question, we screened 22 individual proteins of SARS-CoV-2 for potential ISGylation upon ectopic co-expression of V5-tagged ISG15 and the ISGylation machinery (i.e., E1, E2 and E3) (**Fig. 1A** and **Supplementary Figures 1A,B**). This showed detectable ISGylation for N, NSP7, NSP8, NSP10, NSP12 and NSP16, while the other 16 viral proteins did not undergo ISGylation under these conditions. Of note, the N protein was the most highly ISGylated protein among the SARS-CoV-2 proteins tested where its ISGylation was detectable not only in the ISG15 (V5) blot, but also as ∼15 kDa slower-migrating band in the FLAG blot directly assaying FLAG-tagged N (**Fig. 1A**). As the N protein is actively recruited to the RTC at viral replication organelles where NSP3 resides, and because NSP3’s papain-like protease (PLpro) harbors de-ISGylating activity (35, 45, 48), we next investigated whether PLpro can remove N ISGylation. Ectopic expression of wildtype (WT) SARS-CoV-2 PLpro, but not its catalytically inactive mutant (C111A), abolished the ISGylation of co-expressed N (**Fig. 1B**). Of note, overexpressed PLpro WT, but not C111A mutant, also ablated the ISGylation of co-expressed NSP7, NSP8, NSP10, NSP12 and NSP16 under these conditions (**Fig. 1B** and **Supplementary Figure 1C**). Importantly, N ISGylation was also readily detectable during authentic SARS-CoV-2 infection in A549-hACE2 cells where PLpro’s activity was inhibited using the pharmacological inhibitor GRL-0617 (**Fig. 1C**). Collectively, these results show that the SARS-CoV-2 N protein is modified by ISG15, and that the ISGylation levels of N are controlled by direct de-ISGylation by NSP3’s PLpro enzymatic activity.

**Fig 1.**
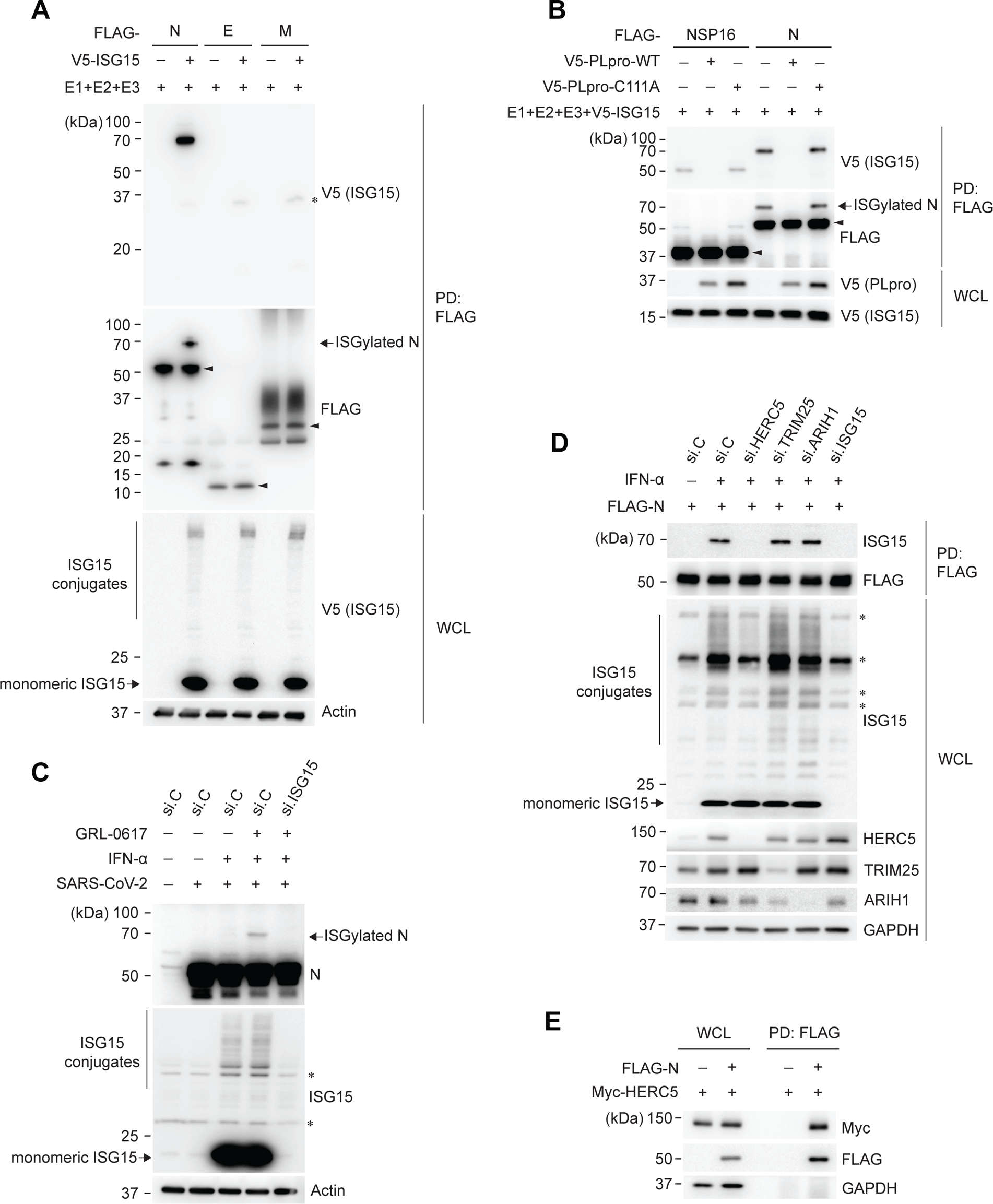
The SARS-CoV-2 N protein is ISGylated by HERC5. (**A**) ISGylation of FLAG-tagged N, E, and M of SARS-CoV-2 in HEK293T cells that were transfected for 24 h with either empty vector or V5-tagged ISG15 along with plasmids encoding E1 (UBE1L), E2 (UBCH8), and E3 (HERC5), determined by FLAG pull-down (PD:FLAG) and immunoblotting (IB) with anti-V5 and anti-FLAG. Whole cell lysates (WCLs) were immunoblotted with the indicated antibodies. (**B**) ISGylation of FLAG-tagged NSP16 and N in HEK293T cells that were co-transfected for 24 h with empty vector or V5-tagged PLpro WT or C111A together with E1, E2, E3, and V5-ISG15, determined by PD:FLAG and IB with anti-V5 and anti-FLAG. (**C**) ISGylation of N protein in A549-hACE2 cells that were transfected for 24 h with either control non-targeting siRNA (si.C) or ISG15-specific siRNA (si.ISG15), followed by mock treatment or stimulation with 1000 U/mL IFN-α for 24 h and subsequent infection with SARS-CoV-2 (MOI 1) for 24 h in the absence or presence of the PLpro inhibitor GRL-0617 (25 µM), determined in the WCLs by IB with anti-N (SARS-CoV-2). WCLs were further immunoblotted with the indicated antibodies. (**D**) ISGylation of FLAG-tagged N in A549 cells transfected with the indicated siRNAs for 24 h, followed by mock treatment or stimulation with 1000 U/mL IFN-α for 24 h and subsequent transfection with FLAG-N for 24 h, determined by PD:FLAG and IB with the indicated antibodies. (**E**) Binding of Myc-tagged HERC5 to FLAG-tagged N in transiently transfected HEK293T cells, determined by PD:FLAG and IB with anti-Myc and anti-FLAG. si.C, nontargeting control siRNA. Asterisks (*) in (**A**), (**C**) and (**D**) indicate non-specific bands. Arrowheads in (**A**) and (**B**) indicate non-ISGylated form of transfected proteins. Data are representative of at least two independent experiments.

### HERC5 is a primary cellular E3 ligase mediating SARS-CoV-2 N ISGylation

HERC5 is a major E3 ligase that ISGylates a variety of host proteins (49). In addition, HERC5 has been shown to ISGylate certain viral proteins (for example, influenza A virus NS1, influenza B virus nucleoprotein (IBV NP), and human papilloma virus L1), which can lead to virus restriction (50–53). As such, we examined the impact of silencing endogenous HERC5 on N ISGylation (**Fig. 1D**). In parallel, we also silenced TRIM25/EFP and ARIH1, which are two other mammalian E3 ligases known to have ISGylating activity (31, 42, 54). Depletion of endogenous HERC5 or ISG15 (the latter serving as a control in this assay) completely abrogated N ISGylation; in contrast, knockdown of endogenous TRIM25 or ARIH1 had no effect on the ISGylation levels of N (**Fig. 1D**). Co-immunoprecipitation (co-IP) assay showed that Myc-tagged HERC5 interacted with FLAG-tagged N (**Fig. 1E**). Together, these data indicate that HERC5 is a primary host E3 ligase that is recruited to and ISGylates the SARS-CoV-2 N protein.

### Identification of the ISGylation sites in SARS-CoV-2 N by mass spectrometry

To identify the specific ISGylation sites in the N protein of SARS-CoV-2, we applied unbiased mass spectrometry (MS) analysis on affinity-purified, ectopically expressed N protein. This identified four lysine residues in N that were covalently linked to ISG15: K266, K355, K387 and K388 (**Fig. 2A,C** and **Supplementary Figure 2A**). Sequence alignment showed that these four lysine residues are conserved in the N proteins of different SARS-CoV-2 variants (**Fig. 2B**). The coronaviral N protein comprises an N-terminal domain (NTD) harboring RNA-binding activity and a C-terminal domain (CTD) required for both RNA binding and N dimerization (7, 8, 55, 56). In addition, N comprises three intrinsically disordered regions (IDRs): the N-IDR, the linker (LKR), and the C-IDR (**Fig. 2A,D**). K266 and K355 are located in the CTD, whereas K387 and K388 localize to the C-IDR that is involved in N oligomerization and phase separation (8, 56) (**Fig. 2A,D**). To validate our MS results, we mutated the four identified sites either individually or in combination, and then compared the ISGylation of these mutant N proteins to that of WT N. This showed that individual mutation of any of the four lysines led to only a slight reduction of N ISGylation, while the combined mutation of K355, K387 and K388 (K355,387,388A) or of all four residues (K266,355,387,388A; hereafter called ‘4KA’ mutant) strongly diminished and near-abolished ISGylation, respectively (**Fig. 2E**). Similarly to the 4KA mutant, a 4KR mutant in which the four ISGylated lysine residues in SARS-CoV-2 N were mutated to arginines, exhibited a loss of ISGylation (as compared to WT N) in cells in which endogenous ISGylation was induced by IFN-α stimulation (**Supplementary Figure 2B**). These results indicate that the SARS-CoV-2 N protein undergoes ISGylation at four residues located in the CTD and the C-IDR. Furthermore, as we consistently observed only one band for ISGylated N (both during authentic SARS-CoV-2 infection and in ectopic N protein expression conditions), this suggests that the N protein is primarily ISGylated at any one of the four sites, and that ISG15 conjugation at one site likely sterically hinders the subsequent ISGylation of additional sites.

**Fig 2.**
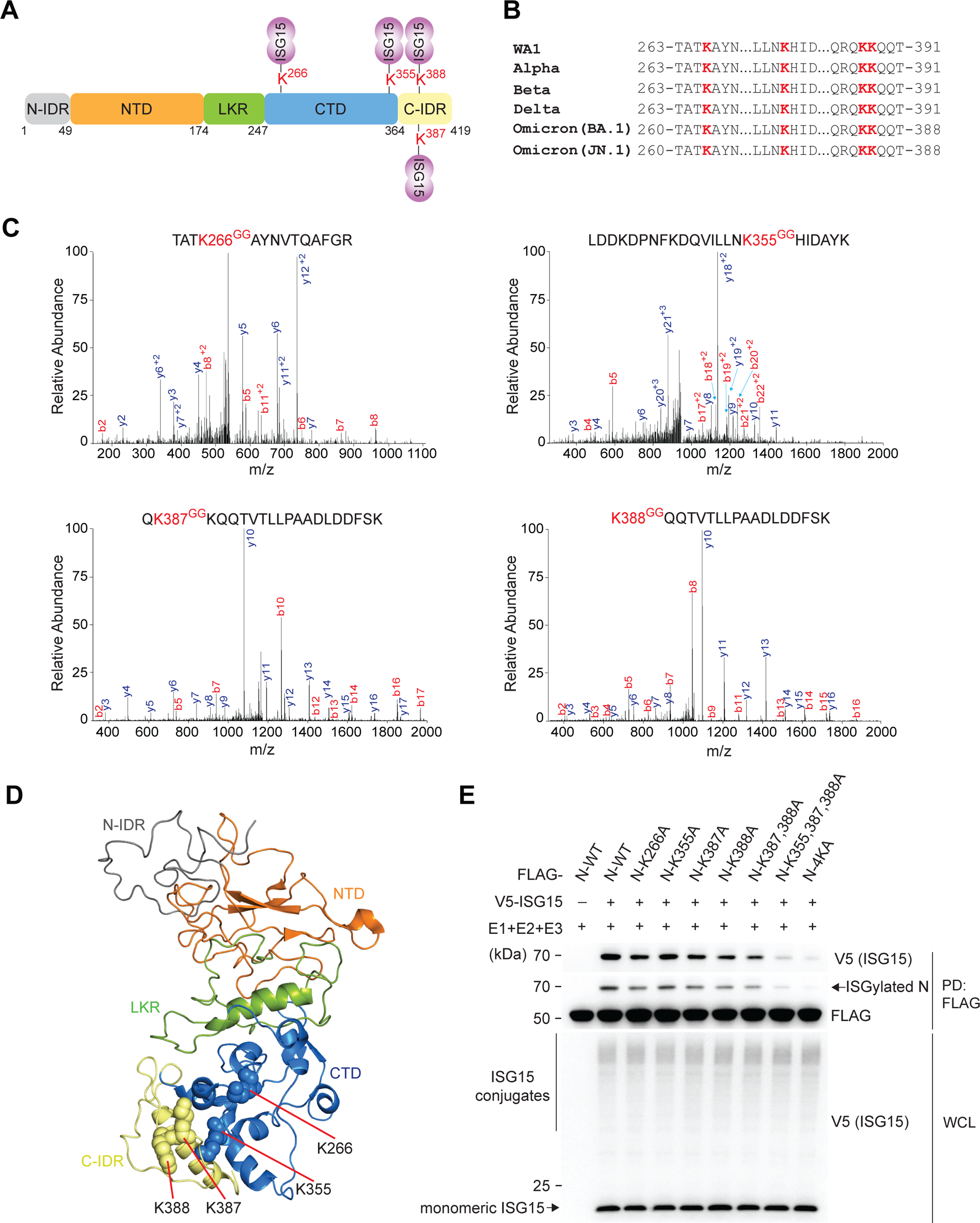
Identification of the ISGylation sites in the SARS-CoV-2 N protein by mass spectrometry. (**A**) Schematic diagram of the domain organization of the SARS-CoV-2 N protein with the ISGylated lysines (Ks) identified by MS analysis illustrated in red. The three intrinsically disordered regions, including the N-IDR, the linker (LKR) and C-IDR, as well as the N-terminal domain (NTD) and C-terminal domain (CTD) are illustrated. (**B**) Amino acid sequence spanning the region of the N protein that contains K266, K355, K387 and K388 for the indicated SARS-CoV-2 variants. The four lysine residues found to be conjugated with ISG15 are highlighted in red. (**C**) Mass spectra of the tryptic peptides of affinity-purified FLAG-tagged N that identified K-GlyGly modifications at K266, K355, K387, and K388. b-and y-ion designations are shown. See also Supplementary Fig. 2A. (**D**) Location of K266, K355, K387, and K388 in the monomeric Cryo-EM structure of full-length N protein (PDB: 8FD5). (**E**) ISGylation of FLAG-tagged N WT or mutants in transiently transfected HEK293T cells that were co-transfected for 24 h with either empty vector or V5-tagged ISG15 together with E1, E2 and E3, determined by PD:FLAG and IB with anti-V5 and anti-FLAG. WCLs were immunoblotted with anti-V5. Data in (**A and C**) are from one unbiased MS screen. Data in (**E**) are representative of at least two independent experiments.

### Ablation of N ISGylation in the context of a recombinant SARS-CoV-2 replicon allows for escape from ISGylation-mediated virus restriction

To determine the physiological role of N ISGylation in virus replication and to obviate any potential gain of function, we developed a single-cycle SARS-CoV-2 replicon (SARS-CoV-2^Rep^) expressing a mScarlet reporter in place of the spike gene, in which mScarlet expression serves as a proxy for viral subgenomic RNA synthesis (**see Methods**). We then mutated the four ISGylated lysine residues in N to arginines (4KR), and then compared the replication of the 4KR and parental (WT) replicons by fluorescence microscopy analyzing mScarlet signals and by qRT-PCR measuring subgenomic and genomic RNA (**Fig. 3A,B**). The SARS-CoV-2 WT and 4KR replicons replicated equally in empty vector-transfected HEK293T cells as well as in cells ectopically expressing the E1-E2-E3 ISGylation machinery. However, in cells that ectopically co-expressed ISG15 WT, we observed a strong inhibition of viral RNA synthesis for the WT replicon, but not the 4KR replicon (**Fig. 3A,B**). Crucially, this difference in ISGylation-mediated virus restriction was not observed in cells that ectopically expressed unconjugatable ISG15-AA (a mutant ISG15 protein where the two glycines required for conjugation were substituted with alanines (35)) (**Fig. 3A,B**), further strengthening that the effects observed for SARS-CoV-2 restriction is due to covalent ISG15 conjugation of the N protein. In line with these data, we detected ISGylation of N from the WT replicon, but not the 4KR replicon, in cells expressing the ISGylation machinery together with ISG15 WT but not ISG15-AA (**Fig. 3C**). These data indicate that ablation of N ISGylation in the context of a recombinant SARS-CoV-2 replicon allows for escape from ISGylation-mediated virus restriction and thus leads to more effective virus replication.

**Fig 3.**
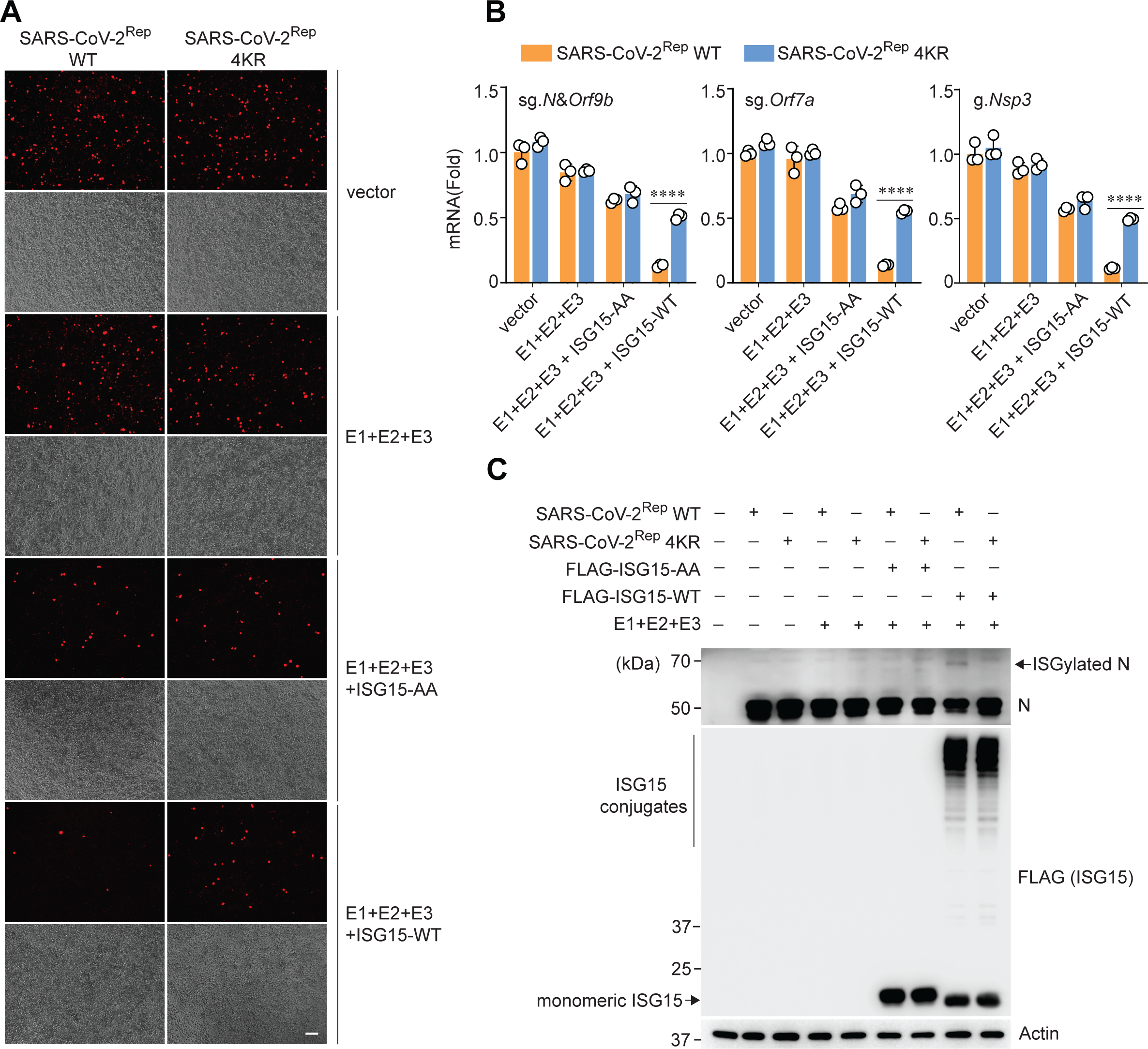
A SARS-CoV-2 replicon encoding an ISGylation-defective mutant N protein escapes HERC5-mediated virus restriction. (**A and B**) Replication of a SARS-CoV-2 replicon encoding either N WT or a mutant in which the four ISGylation sites were mutated to arginine (SARS-CoV-2^Rep^ WT or 4KR), determined by fluorescence microscopy analysis of mScarlet signals (**A**) and qPCR analysis of viral subgenomic and genomic RNA (**B**) in HEK293T cells that were transfected for 48 h with the WT or mutant replicon and empty vector or E1, E2 and E3/HERC5 with or without FLAG-tagged ISG15-WT or ISG15-AA. Scale bar, 100 μm. (**C**) ISGylation of N protein in HEK293T cells that were transfected as in (B), determined in the WCLs by IB with anti-N (SARS-CoV-2). WCLs were further immunoblotted with anti-FLAG (to confirm the expression of FLAG-tagged ISG15 WT or AA) and anti-Actin (loading control). Data are representative of at least two independent experiments (mean ± SD of *n* = 3 biological replicates in (B)). \*\*\*\**P* < 0.0001 (two-tailed, unpaired Student’s *t*-test).

### N ISGylation impedes the oligomerization of N but does not affect its binding to NSP3

The four ISGylation sites localize to the CTD and the C-IDR regions of the SARS-CoV-2 N protein, which have been reported to mediate N oligomerization (19, 56–60). As such, we examined whether N ISGylation affects its ability to form higher-order assemblies by performing co-IP assay assessing the binding of FLAG-tagged N to HA-tagged N. This analysis showed that whereas non-ISGylated N strongly bound to FLAG-tagged N, ISGylated N had diminished binding (**Fig. 4A**). Semi-denaturing detergent agarose gel electrophoresis (SDD-AGE) confirmed that the oligomerization of FLAG-tagged N WT, but not 4KR, is inhibited in the presence of the ISGylation machinery (**Fig. 4B**). As coronaviral N is known to bind to the Ubl1 domain of NSP3 and since this interaction is pivotal for the onset of viral RNA synthesis (61, 62), we also determined whether N ISGylation affects NSP3 binding. Whereas the N protein lost binding to NSP3-ΔUbl1 as expected, both the unmodified and ISGylated N efficiently bound to NSP3-C857A in which the PLpro catalytic activity is ablated (**Supplementary Figure 3A**). These results suggest that N ISGylation does not impede the recruitment of N to NSP3 at viral RTCs but impacts a critical step of viral RNA synthesis thereafter that requires N oligomerization.

**Fig 4.**
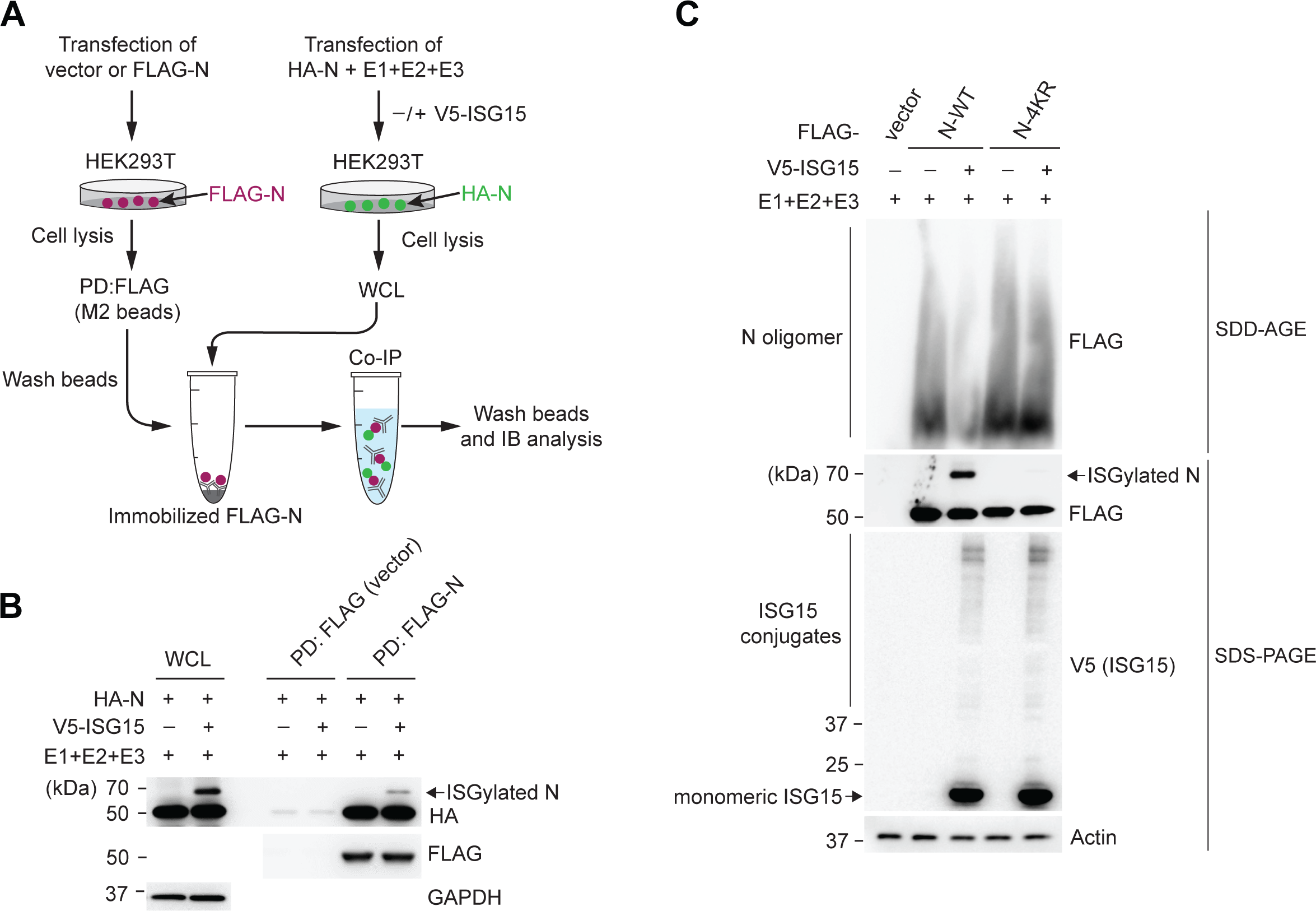
ISGylation of the SARS-CoV-2 N protein impedes its oligomerization. (**A**) Schematic diagram of the co-IP approach to determine the binding of ISGylated vs. non-ISGylated HA-tagged N with FLAG-tagged N. (**B**) Left: FLAG-tagged N (or empty vector as a control) was transfected into HEK293T cells, followed by affinity purification and immobilization using PD:FLAG. Right: In parallel, HA-tagged N was co-transfected with either empty vector or V5-tagged ISG15, together with E1, E2, and E3/HERC5, into HEK293T cells. WCLs were prepared and incubated in vitro with immobilized FLAG-N. Binding was determined by IB with anti-HA and anti-FLAG. (**C**) Oligomerization of FLAG-tagged N WT or 4KR in HEK293T cells that were co-transfected for 24 h with either empty vector or V5-tagged ISG15 together with E1, E2, and E3/HERC5, determined by SDD-AGE and IB with anti-FLAG. WCLs were analyzed by SDS-PAGE and probed by IB with anti-FLAG, anti-V5 and anti-Actin (loading control). Data in (**B**) and (**C**) are representative of at least two independent experiments.

### HERC5 preferentially targets the phosphorylated N protein for ISGylation

The coronaviral N protein is known to undergo phosphorylation at multiple sites, and N phosphorylation states critically regulate its respective functions in the viral life cycle (18, 19, 24, 60, 63–66). Whereas phosphorylated N protein has been reported to promote longer subgenomic RNA synthesis as part of the RTC at early stages of infection, N in its unphosphorylated state drives genome packaging later during infection. Within the highly phosphorylated ‘serine-arginine’ (SR) region located in the LKR of N, two phosphorylation priming sites (S188 and S206) play key roles in N regulation (8, 24). Since PTMs are known to inter-regulate each other (67, 68), we asked whether phosphorylation of the N protein at these sites impacts N ISGylation. The N S188,206A mutant, which has abolished phosphorylation at these sites (evidenced also by a faster-migrating band for N in SDS-PAGE) showed strongly diminished ISGylation compared to WT N (**Fig. 5A**). In accord, treatment of cells with kenpaullone, a pharmacological inhibitor of glycogen synthase kinase 3β (GSK-3β), which is a primary kinase for coronaviral N phosphorylation (24, 25, 66, 69–71), reduced N ISGylation in a dose-dependent manner (**Fig. 5B**). Furthermore, a phosphomimetic mutant of N in which 10 key residues (9 serines and one threonine) in the SR region were mutated to negatively charged aspartic acid residues (termed ‘10D’) (18), showed strongly enhanced ISGylation levels as compared to WT N and the S188,206A mutant (**Fig. 5C**). We posited that the phosphorylation state of N may regulate its interaction with HERC5. Therefore, we next compared the binding of FLAG-tagged N S188,206A (unphosphorylated) and 10D (phosphomimetic) to Myc-HERC5 by co-IP assay and found that the 10D mutant exhibited markedly enhanced HERC5 binding as compared to the S188,206A mutant (**Fig. 5D**). Of note, the 10D mutant also bound to NSP3 more efficiently than N S188,206A did (**Supplementary Figure 3B**). Altogether, these results reveal an intricate crosstalk between N phosphorylation and ISGylation where phosphorylation promotes the recruitment of HERC5 to N and thereby its ISGylation.

**Fig 5.**
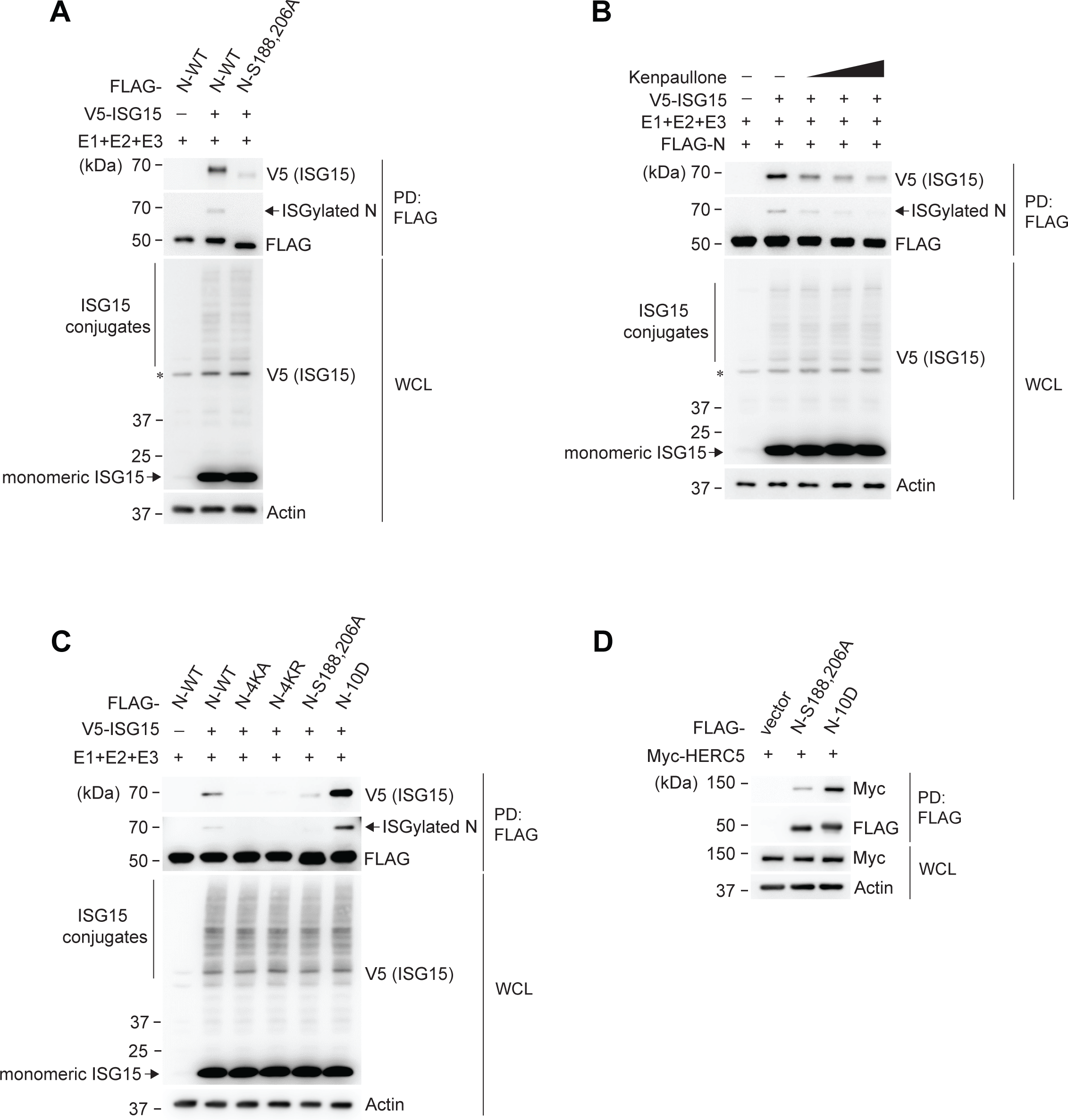
Phosphorylation of the SARS-CoV-2 N protein promotes its ISGylation. (**A**) ISGylation of FLAG-tagged N WT or S188,206A in HEK293T cells that were co-transfected for 24 h with either empty vector or V5-ISG15 along with E1, E2, and E3/HERC5, determined by PD:FLAG and IB with anti-V5 and anti-FLAG. (**B**) ISGylation of FLAG-tagged N in HEK293T cells that were co-transfected for 24 h with either empty vector or V5-ISG15 along with E1, E2, and E3/HERC5 in the absence or presence of the GSK3 inhibitor kenpaullone (5 µM, 10 µM, or 50 µM), determined by PD:FLAG and IB with anti-V5 and anti-FLAG. (**C**) ISGylation of FLAG-tagged N WT or the indicated mutants in HEK293T cells that were co-transfected for 24 h with either empty vector or V5-ISG15 along with E1, E2, and E3/HERC5, determined by PD:FLAG and IB with anti-V5 and anti-FLAG. (**D**) Binding of Myc-tagged HERC5 to FLAG-tagged N S188,206A or 10D in transiently transfected HEK293T cells, determined by PD:FLAG and IB with anti-Myc and anti-FLAG. Asterisks (*) in (**A**) and (**B**) indicate non-specific bands. Data are representative of at least two independent experiments.

## Discussion

ISG15 plays diverse roles in cellular processes, and its covalent conjugation to protein substrates represents a pivotal PTM that orchestrates host antiviral defense (39, 41). Despite the expanding array of host and viral proteins modified by ISG15, *bona fide* protein substrates and their functional alteration post-ISGylation remain poorly characterized. In this study, we screened a near-complete suite of viral proteins from SARS-CoV-2 for ISGylation and identified the N protein as a *bona fide* target. Our work elucidated four specific sites within the N protein that were robustly ISGylated by the E3 ligase HERC5, which impeded N oligomerization and its function in viral RNA synthesis in a manner that is contingent upon its phosphorylation state. As a viral countermeasure, the SARS-CoV-2 PLpro actively removed ISG15 from the conjugated N protein via de-ISGylating activity, which is expected to reinstate higher-order N assembly facilitating viral replication. These findings expand the repertoire of viral proteins susceptible to direct ISGylation and provide a new paradigm of SARS-CoV-2–host arms race that is regulated by the ISGylation–deISGylation circuit.

The coronaviral N protein is an essential structural component of virus particles and exerts diverse intracellular functions in viral replication (2, 72). The four ISGylation sites identified in SARS-CoV-2 N localize to the CTD (K266 and K355) and the C-IDR (K387 and K388), which have been implicated in protein dimerization, oligomerization, RNA binding, and phase separation (7, 8, 55, 56). In line with studies demonstrating that deletion of the CTD and C-IDR in the SARS-CoV and 229E N proteins abrogated their oligomerization (57, 58), our results showed that ISGylation at the four lysine residues diminished SARS-CoV-2 N oligomerization. Intriguing, while the four ISGylated lysine residues are not directly involved in dimer interface interactions, K266 and K355 in the CTD are proximal to the swapped β-strands of the dimer interface that is postulated to form an RNA binding groove (**Supplementary Figure 4A,B**) (7, 55). It is thus possible that the bulky ISGylation at K266 and K355 may influence N protein conformational dynamics crucial for dimer formation and RNA binding, which seed higher-order oligomeric assembly of N. It is worth noting that the longest dimension of ISG15 (∼15 kDa) is approximately 53 angstroms, nearly equal to the longest dimension of the N CTD dimer (∼26 kDa) (**Supplementary Figure 4C**) (26, 55). Therefore, one ISGylation event nearly doubles the size of the N CTD dimer unit alone. Crosslinking MS analysis of the SARS-CoV-2 N protein also revealed a cluster of lysine residues within the IDR contributing to intermolecular and/or intramolecular N interactions (19, 60). Deletion of the highly crosslinked region encompassing two of the ISGylated residues identified in our study (K387 and K388) did not affect N binding to viral RNA but abolished N condensate formation (i.e. phase separation) (60), suggesting that ISGylation of K387 or K388 in the C-IDR may directly impact oligomeric N assembly. Therefore, even a very small pool (<5%) of ISG15-conjugated N may exert a dominant negative effect on oligomeric assembly and thereby the function of unconjugated N in infected cells due to the bulk steric effect of ISGylation. Future structural analyses of the SARS-CoV-2 N protein in complex with ISG15 will be required to unveil the precise mechanism(s) of how ISGylation modulates N oligomerization and its functions.

Our MS analysis identified K266, K355, K387, and K388 as specific ISGylation sites; however, we consistently observed only one band for ISGylated N, both in vitro and during authentic SARS-CoV-2 infection. One explanation for this could be that ISGylation at any of these four lysine residues may hinder the ISGylation machinery’s access to the other sites, akin to what has been observed for IAV NS1 where ISG15 conjugation to any lysine within a hotspot consisting of multiple nearby ISGylatable lysines was mutually exclusive and gave rise to only a single band of the same mobility on SDS-PAGE gels (73). Considering the N protein’s dynamic interactions with various host proteins (74, 75), it is also possible that some of these interactions could mask some of the ISGylation sites. Structural studies of ISGylated N are needed to elucidate the molecular details of how ISGylation at one specific site impacts the ISGylation of the other sites. Future studies will also need to determine the precise stoichiometry of the ISGylated vs. non-ISGylated pool of N during SARS-CoV-2 infection as well as the spatial and temporal aspects of its regulation by HERC5 and viral or host deISGylating enzymes (i.e., PLpro and USP18). Lastly, although our studies utilizing unbiased MS analysis and biochemical approaches revealed four major ISGylation sites for N, we cannot rule out that there are additional sites in N that may undergo ISGylation.

Inhibition of SARS-CoV-2 N oligomerization by ISGylation is reminiscent of the dominant-negative effect that ISGylation has on IBV NP oligomerization, which ultimately inhibits the assembly of the viral ribonucleoprotein complex (51). While the inhibitory role of ISGylation on NP function is counteracted through viral NS1-mediated sequestration of ISGylated NP, coronaviral PLpro directly removes ISGylation from N. Thus, SARS-CoV-2 via PLpro exerts an effective mechanism to evade ISG15-mediated direct virus restriction; as such, we could detect N ISGylation in SARS-CoV-2-infected cells only after chemical inhibition of the PLpro enzymatic activity. Additionally, SARS-CoV-2 is equipped with a number of IFN antagonists keeping the amount of produced IFNs at relatively low levels, which is expected to limit the activity of the cellular ISG15 conjugation machinery (of note, several of its components are IFN inducible including ISG15 itself) (76–79). Therefore, NSP3 PLpro has evolved a two-pronged mechanism to escape ISGylation-dependent virus restriction (i.e., by evading innate immune activation and thus ISG15 upregulation (35, 45) and via direct deISGylation of N as reported herein).

Multiple PTMs regulate the coronaviral N protein, including phosphorylation, ubiquitination, glycosylation, methylation, acetylation, and SUMOylation (18–21, 23, 63, 64, 80–82). However, it remains obscure how these PTMs functionally orchestrate the distinct activities of N during virus infection. Essential for the SARS-CoV-2 life cycle is phosphorylation of N at specific serine residues within the highly conserved SR region, which is dynamically regulated by several host kinases and phosphatases (24, 70, 71, 83). Different phosphorylation states of N dictate its distinct functions, with the phosphorylated and unphosphorylated pool of N participating in viral RNA synthesis and genome packaging, respectively (65, 66). These functions were later linked to distinct states of phase separation of the N protein regulated by phosphorylation. Phosphorylated N protein forms low-viscosity liquid-like droplets conducive to viral RNA synthesis at the RTC, while unphosphorylated N assembles into high-viscosity condensates essential for viral RNA packaging (18–20, 84). Accordingly, our findings revealing that phosphorylated (but not unphosphorylated) N is robustly ISGylated suggest that the N proteins at the RTC are the primary target of the HERC5 ISGylation machinery. Suppression of oligomeric assembly of phosphorylated N by ISGylation directly inhibits viral RNA synthesis, as evidenced by our data from a single-cycle replicon system that circumvents any accumulative effects from multi-cycle viral replication. This antiviral effect of N ISGylation on viral RNA synthesis therefore sets the SARS-CoV-2 mechanism apart from that of NP ISGylation during IBV infection, which instead targets the formation of viral ribonucleoproteins, a later event of the viral life cycle (51).

Our data also suggested that the viral NSP3 protein selectively bound phosphorylated N protein, underscoring a spatially resolved PLpro antagonism during SARS-CoV-2 infection. DeISGylation of phosphorylated N by PLpro in close proximity at the RTC is conceivably more efficient for the virus to rapidly overcome RNA synthesis blockage than recruiting a cellular deISGylating enzyme, such as USP18. Indeed, USP18 preferentially bound to the ‘phospho-null’ N mutant (S188,206A) (**Supplementary Figure 3C**), suggesting that USP18-mediated deISGylation may be co-opted by the virus to serve as a secondary safeguard mechanism against misincorporation of ISGylated N during RNA packaging. Further research is warranted to uncover the molecular details of how viral and host deISGylating enzymes regulate N function at different stages of the SARS-CoV-2 life cycle. Along these lines, elucidating the PTM regulatory network governing the crucial functions of the N proteins of SARS-CoV-2 and other coronaviruses is vital for comprehending coronavirus biology and may pave the way for developing new antiviral therapies.

## Acknowledgements

We thank Bellinda Willard (Proteomics Core of Cleveland Clinic’s Lerner Research Institute) for support with mass spectrometry analysis. We are grateful to Cindy Chiang (Cleveland Clinic Florida Research and Innovation Center) for her support with MS data visualization. This study was funded by the US National Institutes of Health grants R37 AI087846 and DP1 AI169444 (to M.U.G.). The Fusion Lumos instrument used for mass spectrometry analysis was purchased via an NIH shared instrument grant 1S10OD023436-01.

## Author Contributions

Conceptualization: J.Z., G.L., M.U.G.; Formal analysis: J.Z., G.L.; Funding acquisition: M.U.G.; Investigation: J.Z., G.L. and C.M.G., with J.Z. performing all experiments; G.L. providing key DNA constructs and cell lines and establishing the SARS-CoV-2 replicon system; and C.M.G. performing analysis of PDB structures. Supervision: S.R.S, M.U.G.; Visualization: J.Z., C.M.G.; Writing original draft: J.Z., G.L., M.U.G.

## Materials and Methods

### Cell culture and transfections

HEK293T and A549 cells were purchased from American Type Culture Collection (ATCC) and maintained in Dulbecco’s modified Eagle’s medium (DMEM, Gibco) supplemented with 10% (v/v) fetal bovine serum (FBS, Gibco), 1 mM sodium pyruvate (Gibco) and 100 U/mL of penicillin–streptomycin (Gibco). Vero E6-TMPRSS2 cells were maintained in DMEM supplemented with 10% (v/v) FBS, 1 mM sodium pyruvate, 100 U/mL of penicillin– streptomycin and 40 μg/mL blasticidin (Invivogen). HEK293T and A549 cells inducibly expressing human ACE2 were generated by lentiviral transduction (pGLVX-TetOne-hACE2) followed by selection with puromycin (1 μg/mL and 2 μg/mL, respectively). Transient DNA transfections were performed using linear poly(ethylenimine) (1 mg/mL of solution in 10 mM Tris-HCl, pH 6.8; Polysciences), Lipofectamine 2000 (Invitrogen), or *Trans*IT-X2 Transfection Reagent (Mirus) as per the manufacturers’ instructions.

### Plasmids

FLAG-tagged SARS-CoV-2 constructs (NSP4, NSP14, NSP16, ORF3, ORF6, ORF7a, ORF7b, ORF9b, ORF9c, N, E, M) were kindly provided by Jae U. Jung (Cleveland Clinic Lerner Research Institute). Strep-tagged SARS-CoV-2 constructs (NSP1, NSP2, NSP5, NSP7, NSP8, NSP9, NSP10, NSP11, NSP12, NSP15) were described previously (85). pCAGGS-HA-Ube1L, pFLAG-CMV2-UbcH8, pcDNA-3×Myc-HERC5, pCAGGS-V5-ISG15, pcDNA-3×FLAG-ISG15, pcDNA-3×FLAG-ISG15 (GG156/157AA), pcDNA-V5-PLpro, and pcDNA-V5-PLpro (C111A) have been described previously (33, 35). The WT and ISGylation-site mutants of N (K266A; K355A; K387A; K388A; K387,388A; K355,387,388A; 4KA; and 4KR) and phosphorylation mutants of N (S188,206A; and 10D) were generated by site-directed mutagenesis and were subcloned into pcDNA3.1(-) harboring a C-terminal 3×FLAG tag. HA-tagged N was generated by amplifying the N ORF using FLAG-tagged N as a template and then subcloned into pcDNA3.1(-) containing an N-terminal 3×HA tag. pcDNA-V5-NSP3 was generated by amplifying the full-length NSP3 ORF from pDONR207 SARS-CoV-2 NSP3 (Addgene #141257; a gift from Fritz Roth (86)) and by subcloning it into the pcDNA3-C-V5 vector. The C857A and ΔUbl1 mutants of NSP3 were generated by site-directed mutagenesis and overlap extension PCR. pCMV-HA-USP18 was generated through subcloning using Flag-HA-USP18 (Addgene #22572; a gift from Wade Harper (87)) as a template.

### Viruses

SARS-CoV-2 (strain K49) was rescued from a bacterial artificial chromosome (BAC) encoding hCoV-19/Germany/BY-pBSCoV2-K49/2020 (GISAID EPI_ISL_2732373), which was kindly provided by Armin Ensser (Friedrich-Alexander University Erlangen-Nürnberg, Germany) (88), and was propagated in Vero E6-TMPRSS2 cells (89). All work relating to SARS-CoV-2 live virus was conducted in the BSL-3 facility of the Cleveland Clinic Florida Research and Innovation Center (CC-FRIC). All work was reviewed and approved by the CC-FRIC Institutional Biosafety Committee in accordance with U.S. National Institutes of Health guidelines.

### Co-Immunoprecipitation and immunoblot analysis

Cells that were transfected as indicated were lysed in 1% Nonidet P-40 buffer (50 mM HEPES pH 7.4, 150 mM NaCl, 1% (v/v) NP-40, 1 mM EDTA, 1× protease inhibitor cocktail (Sigma, Cat# P2714) and cleared by centrifugation at 21,000 ×*g* and 4°C for 15 min. Cell lysates were then subjected to FLAG pulldown (PD) using anti-FLAG (M2) magnetic beads or anti-FLAG M2 agarose beads (MilliporeSigma) at 4°C for 4-16 h. For IP with anti-V5, cell lysates were incubated with Protein G Dynabeads conjugated with the anti-V5 antibody at 4°C for 4-16 h. The beads were washed with NP-40 buffer and, afterwards, the proteins were eluted by heating in 1× Laemmli SDS sample buffer at 95°C for 5 min. Protein samples were resolved on Bis–Tris SDS-polyacrylamide gel electrophoresis (PAGE) gels, transferred on to polyvinylidene difluoride (PVDF) membranes (Bio-Rad), and visualized using the SuperSignal West Pico PLUS or Femto chemiluminescence reagents (Thermo Fisher Scientific) on an ImageQuant LAS 4000 Chemiluminescent Image Analyzer (General Electric) as previously described (35).

### Antibodies and other reagents

Primary antibodies used in the present study include anti-ISG15 (1:500, Cat# sc-166755; Santa Cruz), anti-ISG15 (1:1000, Cat# 15981-1-AP; Proteintech), anti-NSP3 (1:1,000, Cat# GTX135589; GeneTex), anti-Nucleocapsid (N) (1:1,000, Cat# 33717; CST), anti-FLAG Rabbit mAb (1:1,000, Cat# 14793; CST), anti-HA (1:1,000, Cat# 3724; CST), anti-V5 (1:1,000, Cat# R960-25; Invitrogen; used for IP), anti-V5 (1:1,000, Cat# 13202; CST; used for immunoblotting of WCLs), anti-HERC5 (1:1,000, Cat# 703675; Invitrogen), anti-TRIM25/EFP (1:1,000, Cat# 610570; BD Biosciences), anti-ARIH1 (1:500, Cat# sc-390763; Santa Cruz), anti-Actin (1:2,000, Cat# GTX629630; GeneTex), anti-GAPDH (1:1,000, Cat# 97166; CST), anti-Myc (1:1,000, Cat# 2276; CST), and anti-Strep (1:1,000, Cat# NC9261069, IBA Lifesciences). Anti-FLAG M2 magnetic beads (Sigma-Aldrich, Cat# M8823), anti-FLAG M2 agarose beads (Sigma-Aldrich, Cat# A2220), and Strep-Tactin® Sepharose® resin (IBA Lifesciences, Cat# 2-1201-010) were used for pull-down assays. GRL-0617 was purchased from AdooQ Bioscience. Universal type I IFN (human IFN-alpha hybrid protein) was obtained from PBL Assay Science (Cat# 11200–2).

### Semi-denaturating detergent agarose gel electrophoresis (SDD-AGE)

SARS-CoV-2 N protein oligomerization was examined by SDD-AGE using a previously described protocol (35) with slight changes. Briefly, HEK293T cells transfected for 24 h with FLAG-tagged N were lysed in a buffer containing 50 mM HEPES (pH 7.4), 150 mM NaCl, 0.5% (v/v) NP-40, 10% (v/v) glycerol and 1× protease inhibitor cocktail (MilliporeSigma) at 4°C for 20 min. Cell lysates were cleared by centrifugation at 16,000 ×*g* and 4°C for 10 min and then incubated on ice for 1 h. Cell lysates were then combined with 4× SDD-AGE buffer (2× Tris/borate/EDTA (TBE), 40% (v/v) glycerol, and 8% (w/v) SDS) to reach 1× (0.5× Tris/borate/EDTA (TBE), 10% (v/v) glycerol, and 2% (w/v) SDS), and then incubated at room temperature for 5 min. Samples were then loaded onto a vertical 1.5% agarose gel containing 1× TBE and 0.1% (w/v) SDS and run at 100V for 40 min at 4°C. Proteins were transferred onto a PVDF membrane and analyzed by IB with anti-FLAG.

### RNA isolation and quantitative reverse transcription-PCR

Total cellular RNA was purified using the E.Z.N.A. HP Total RNA Kit (Omega Bio-tek) following the specific instructions by the manufacturer. One-step RT-qPCR was performed using the SuperScript III Platinum One-Step qRT-PCR Kit (Invitrogen) and custom PrimeTime qPCR Probe Assays (IDT) for SARS-CoV-2 subgenomic RNA (sg.*N&ORF9b*, sg.*ORF7a*) and genomic RNA (NSP3 region; g.*NSP3*) (90) on a QuantStudio 6 Pro Real-Time PCR System (Applied Biosystems). Relative mRNA expression was normalized to the levels of *GAPDH* and expressed relative to the values for control cells using the ΔΔC_t_ method.

### Knockdown mediated by siRNA

Transient knockdown in A549 or A549-hACE2 cells was performed using SMARTpool small interfering (si)RNAs (Horizon Discovery) as previously described (91). The specific siRNAs used in this study are: HERC5 (L-005174-00-0005), TRIM25/EFP (L-006585-00-0005), ARIH1 (L-019984-00-0005), ISG15 (L-004235-03-0005) and Non-targeting Control Pool (D-001810-10-05). Efficient knockdown of the respective proteins was confirmed in the WCLs by IB using specific antibodies.

### SARS-CoV-2 replicon and generation of the 4KR N mutant

The SARS-CoV-2 reporter replicon expressing the full-length genomic RNA of SARS-CoV-2 (strain K49) containing mScarlet3 in place of the spike gene was *de novo* assembled using viral cDNA fragments and a single-copy F plasmid backbone (*repE*– *parABC*). A mini-arabinose operon cassette (*AraC*–*trfA*) was added to induce amplification of plasmid copy number. Briefly, a fragment containing human cytomegalovirus (CMV) enhancer–promoter, 5′ UTR, partial CDS of *NSP1*, *ORF3a– ORF10* CDS, 3′ UTR, hepatitis delta virus ribozyme, and bovine growth hormone polyadenylation signal (bGH polyA) was generated by overlap extension PCR and pre-assembled with the arabinose cassette and the F plasmid backbone fragments devoid of *Bsa*I restriction sites. A linker sequence containing two *Bsa*I sites was inserted between *NSP1* and *ORF3a–ORF10* CDS to facilitate subsequent Golden Gate Assembly. A T7 and a tonB terminator were further introduced to flank the CMV–bGH polyA cassette to prevent deleterious read-through transcription in *E. coli*. The remaining viral genome portion (from partial CDS of *NSP1* to *NSP16*) was then divided into 5 fragments and, along with a synthetic fragment harboring mScarlet3 CDS downstream of the authentic spike gene transcription regulatory sequence (TRS), cloned into the pre-assembled vector using the NEBridge® Golden Gate Assembly Kit (*Bsa*I-HF® v2; NEB). A synonymous point mutation (A17973G) was introduced to the *NSP13* gene to remove an internal *Bsa*I site. The assembled replicon was rescued in NEB® 10-beta Competent *E. coli* (NEB), and the complete sequence was confirmed by Nanopore sequencing (Plasmidsaurus).

The 4KR N mutant replicon was generated by amplifying a mutation-containing fragment via overlap extension PCR and ligating it into the parental WT replicon between *Psp*XI and *Kfl*I. To induce replicon copy number, bacterial cultures grown overnight at 30°C were diluted 1:50 in fresh LB broth, and *L*-arabinose was supplemented to the culture when OD_600_ reached 0.4, at a final concentration of 0.01%. Replicon DNA was isolated after another 6-hour growth using the ZymoPURE™ II Plasmid Midiprep Kit (Zymo Research). Reporter replicon-transfected HEK293T cells were imaged 48 h after transfection on an EVOS M5000 Imaging System (Thermo Fisher Scientific) equipped with an RFP (531/593 nm) light cube and a 10x objective.

### Mass spectrometry analysis of N protein ISGylation

To identify the ISGylation sites of the SARS-CoV-2 N protein, HEK293T cells were transfected in three groups, each consisting of approximately 4 x 10^7^ cells. The first group was transfected with empty vector (26 μg); the second group with FLAG-tagged N (18 μg) together with empty vector (8 μg); and the third group with FLAG-tagged N (18 μg) together with V5-tagged ISG15 (8 μg). Each group was additionally co-transfected with E1 (18 μg), E2 (8 μg), and E3/HERC5 (18 μg) for 24 h. After harvesting, cells were lysed in 1% Nonidet P-40 buffer (50 mM HEPES pH 7.4, 150 mM NaCl, 1% (v/v) NP-40, 1 mM EDTA, 1× protease inhibitor cocktail (Sigma, Cat# P2714), clarified by centrifugation at 21,000 ×*g* for 15 min at 4°C, and then incubated with anti-FLAG (M2) magnetic beads at 4°C for 4 h. Pulldown samples were washed with 1M RIPA buffer (20 mM Tris-HCl pH 8.0, 1M NaCl, 1% (v/v) NP-40, 1% (w/v) deoxycholic acid, 0.01% (w/v) SDS) and afterwards resolved by SDS-PAGE. The protein gel was stained with GelCode Blue Stain Reagent (Thermo Fisher) for 1 h at room temperature and then destained with UltraPure water (Invitrogen) for 4 h at room temperature. The unique band corresponding to the size of ISGylated N protein (around 70 kDa) in the V5-ISG15 overexpressed group was excised and then analyzed by LC-MS/MS at the Proteomics Core of Cleveland Clinic’s Lerner Research Institute (Ohio).

### Structural modeling and sequence alignment

SARS-CoV-2 N structures (PDBs: 8FD5, 7XWX, 7UXX, and 1Z2M) were accessed from the RCSB Protein Data Bank (92), aligned and visualized using PyMOL (Version 2.5.5). Primary amino acid sequence alignments of the N proteins from different SARS-CoV-2 variants were performed using Clustal Omega.

### Statistical analysis

Data were analyzed using GraphPad Prism software (version 10), and statistical analyses were conducted using a two-tailed Student’s *t*-test (unpaired). A p value of < 0.05 was considered statistically significant. In general, three biological replicates for each condition were used, unless indicated otherwise. Data were reproduced in independent experiments as described in the legend for each figure.

**Fig S1. ISGylation of SARS-CoV-2 proteins and their deISGylation by PLpro.**

**(A and B)** ISGylation of the indicated Strep-tagged (A) or FLAG-tagged (B) SARS-CoV-2 proteins in HEK293T cells that were co-transfected for 24 h with either empty vector or V5-tagged ISG15 together with E1, E2, and E3, determined by PD:Strep (A) or PD:FLAG (B) and IB with anti-V5 (A and B) and anti-Strep (A) or anti-FLAG (B). (C) ISGylation of Strep-tagged NSP7, NSP8, NSP10, and NSP12 in HEK293T cells that were co-transfected for 24 h with either empty vector or SARS-CoV-2 PLpro WT or C111A together with E1, E2, E3 and V5-ISG15, determined by PD:Strep and IB with anti-V5 and anti-Strep. Asterisks (*) in (A) and (B) indicate non-specific bands. The red and black arrowheads indicate ISGylated and non-ISGylated forms of the transfected proteins, respectively.

Data are representative of at least two independent experiments.

**Fig S2. Identification and biochemical validation of the ISGylation sites in the SARS-CoV-2 N protein. (A)** LC-MS/MS analysis of ISGylated FLAG-tagged N. For K266, a triply charged peptide with an observed m/z of 547.9491 Da was identified and had a mass within -0.98 ppm of the expected mass for the TATKAYNVTQAFGR + GG peptide. The MS/MS spectra for this peptide contained several C-terminal y and N-terminal b ions and the masses of the y11 and b5 ions are consistent with modification at K266. For K355, a triply charged peptide with an observed m/z of 952.83356 Da was identified and had a mass within -1.02 ppm of the expected mass for the LDDKDPNFKDQVILLNKHIDAYK + GG peptide. The MS/MS spectra for this peptide contained several C-terminal y and N-terminal b ions and the mass difference between the y6 and y7 ions is consistent with modification at K355. For K387, a doubly charged peptide with an observed m/z of 1180.6259 Da was identified and had a mass within -0.4 ppm of the expected mass for the QKKQQTVTLLPAADLDDFSK + GG peptide. The MS/MS spectra for this peptide contained several C-terminal y and N-terminal b ions and the mass of the b2 ion is consistent with modification at K387. For K388, a doubly charged peptide with an observed m/z of 1052.5479 Da was identified and had a mass within - 1.62 ppm of the expected mass for the KQQTVTLLPAADLDDFSK + GG peptide. The MS/MS spectra for this peptide contained several C-terminal y and N-terminal b ions and the mass of the y16 ion is consistent with modification at K388. (B) ISGylation of FLAG-tagged SARS-CoV-2 N WT or the indicated mutants in transiently transfected A549 cells that were pre-treated with IFN-α (1000 U/mL) for 24 h, determined in the WCLs by IB with anti-FLAG. WCLs were further immunoblotted with anti-ISG15 and anti-Actin. Asterisks (*) indicate non-specific bands.

Data in in (A) are from one unbiased MS screen. Data in (B) are representative of at least two independent experiments.

**Fig S3. SARS-CoV-2 N ISGylation does not affect NSP3 binding, and N phosphorylation differentially regulates N binding to NSP3 and USP18. (A)** Binding of FLAG-tagged N to NSP3 WT or its mutants (ΔUbl1 or C857A) in HEK293T cells that were co-transfected for 24 h with either empty vector or FLAG-tagged ISG15 together with E1, E2, and E3/HERC5, determined by IP:V5 and IB with anti-FLAG and anti-V5. (B) Binding of V5-tagged NSP3 to FLAG-tagged N S188,206A or 10D in transiently transfected HEK293T cells, determined by PD:FLAG and IB with anti-V5 and anti-FLAG. (C) Binding of HA-tagged USP18 to FLAG-tagged N S188,206A or 10D in transiently transfected HEK293T cells, determined by PD:FLAG and IB with anti-HA and anti-FLAG.

Data are representative of at least two independent experiments.

**Fig S4. Structural analysis of ISGlyation sites in the N protein.**

**(A)** Alignment of the SARS-CoV-2 N protein CTD dimer structure (monomer 1 in cyan and monomer 2 in dark green, PDB: 7XWX) with the CTD domain from full length monomeric N protein (marine blue, PDB: 8FD5). Locations of K266 and K355 (shown as spheres) are proximal to the swapped β-strands of the dimer interface. (B) K266 and K355 (shown as spheres) mapped onto the SARS-CoV-2 N CTD dimer with vacuum electrostatic potential surface map shown (PDB: 7UXX) and putative RNA binding grove denoted (55). (C) Relative size comparison of ISG15 (salmon, PDB: 1Z2M) and the N protein CTD dimer (blue and green, PDB: 7UXX) with approximate longest dimensions shown.

