## Supplementary figures and images for "ISGylation of the SARS-CoV-2 N protein by HERC5 impedes N oligomerization and thereby viral RNA synthesis"

### Supplementary Figure 1

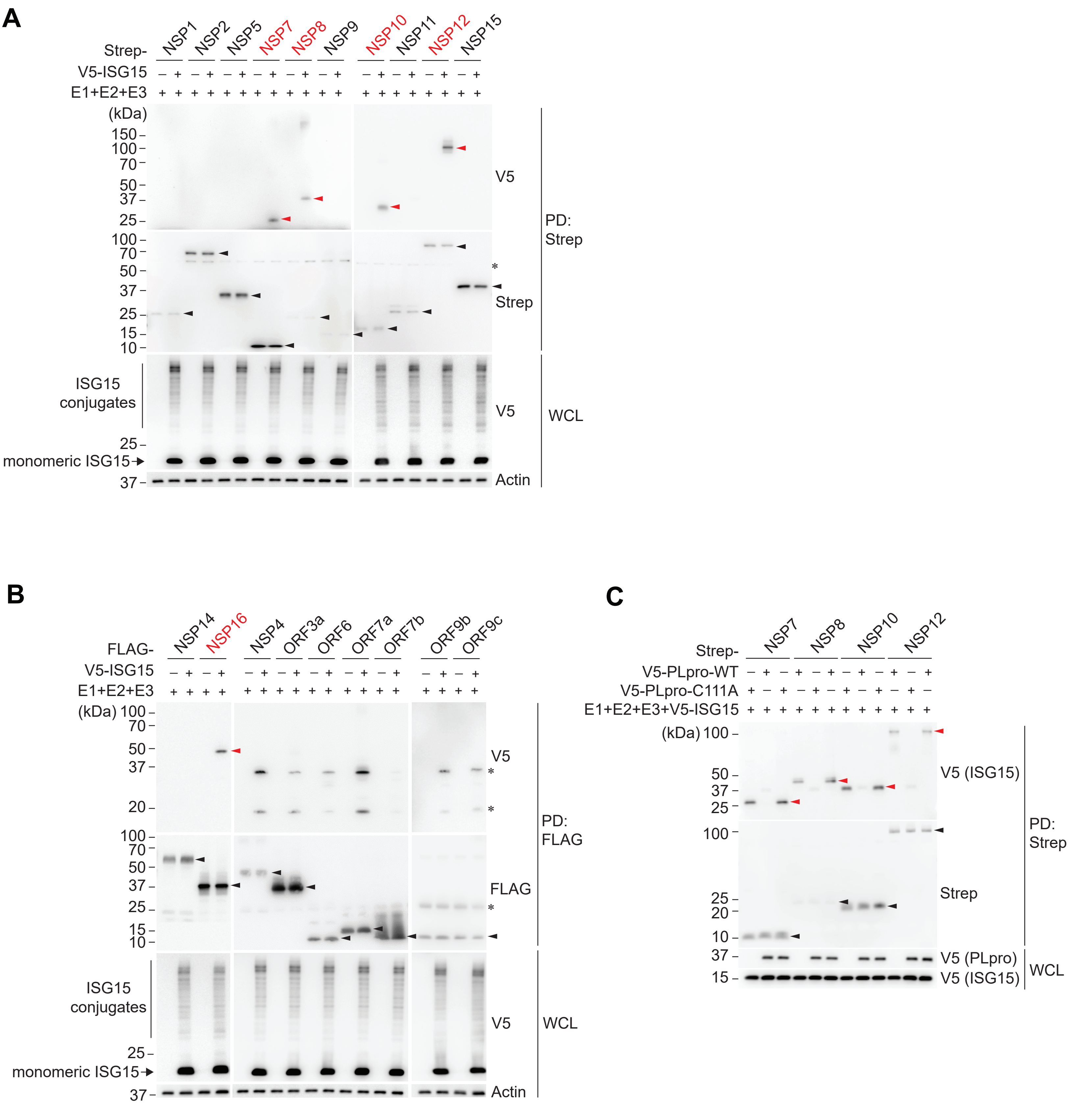


**Supplementary Figure 1**

### Supplementary Figure 3

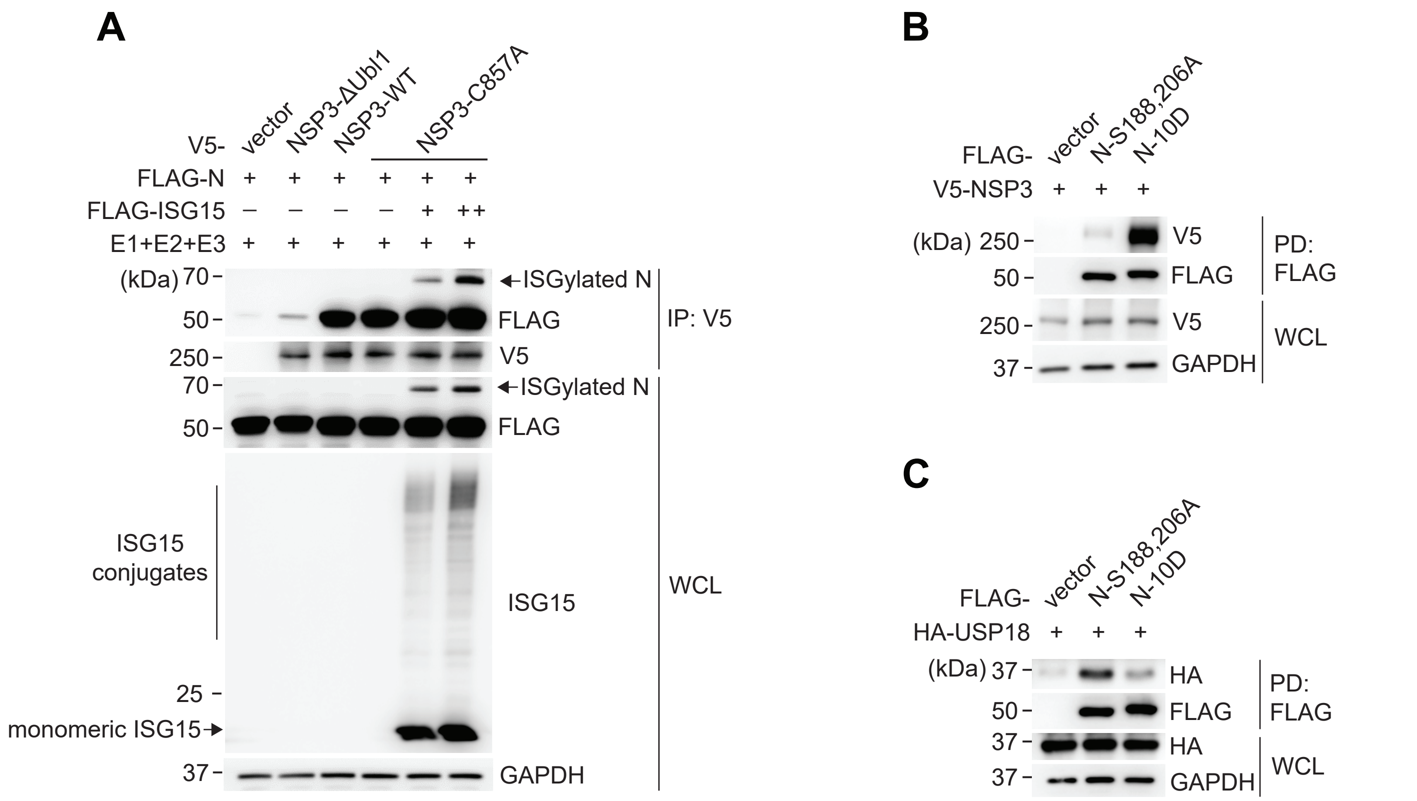


**Supplementary Figure 3**

### Supplementary Figure 4

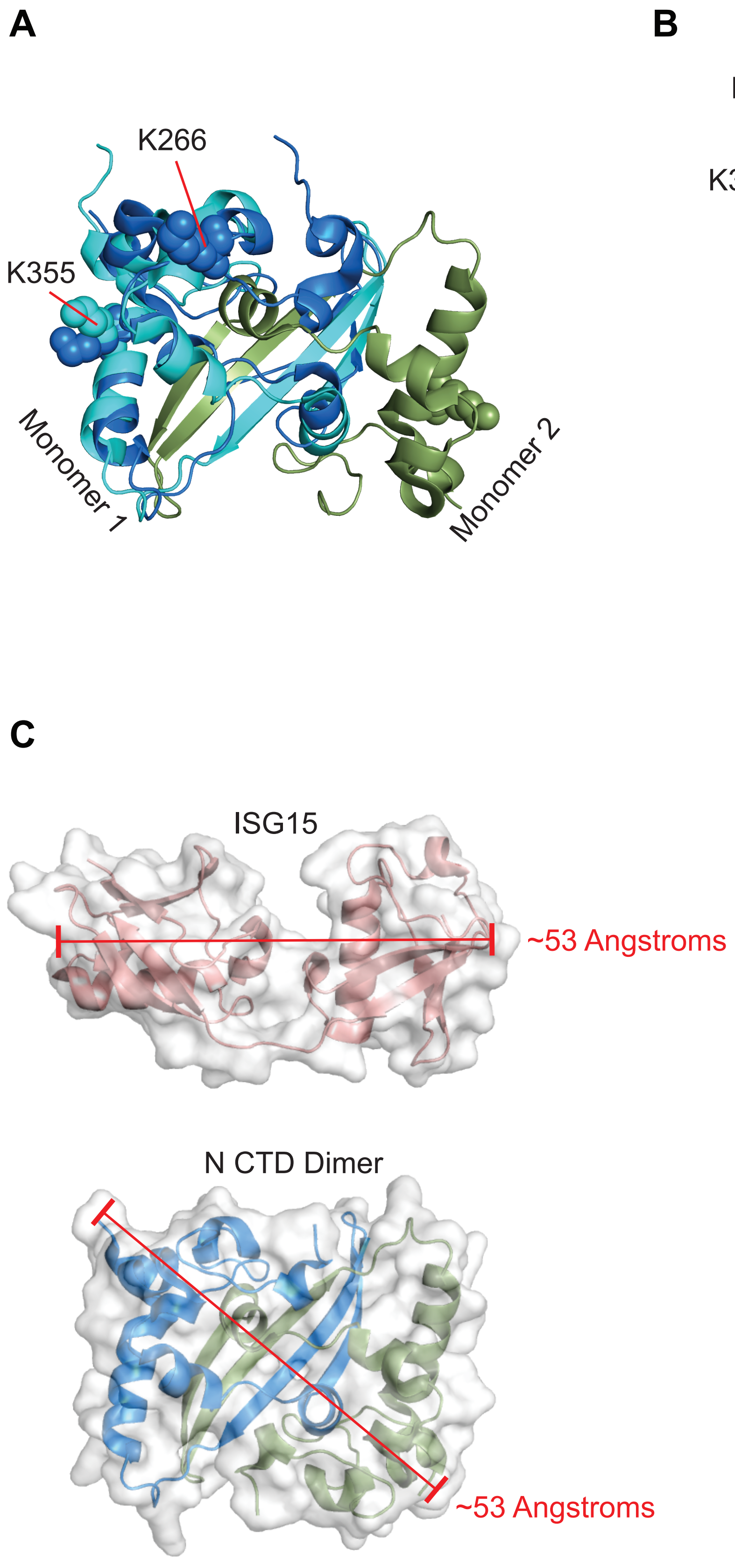


**Supplementary Figure 4**
