## Supplementary Figure 2 for "ISGylation of the SARS-CoV-2 N protein by HERC5 impedes N oligomerization and thereby viral RNA synthesis"

**A**

| Site | [M+H] | m/z | Δm (ppm) | Sequence | XCorr Score |
|---|---|---|---|---|---|
| 266 | 1641.8 | 547.9491 | -0.98 | TATK$^{GG}$AYNVTQAFGR | 3.91 |
| 355 | 2856.5 | 952.83356 | -1.02 | LDDKDPNFKDQVILLNK$^{GG}$HIDAYK | 5.28 |
| 387 | 2360.2 | 1180.62598 | -0.4 | QK$^{GG}$KQQTVTLLPAADLDDFSK | 4.61 |
| 388 | 2104.1 | 1052.54797 | -1.62 | K$^{GG}$QQTVTLLPAADLDDFSK | 6.54 |

**B**

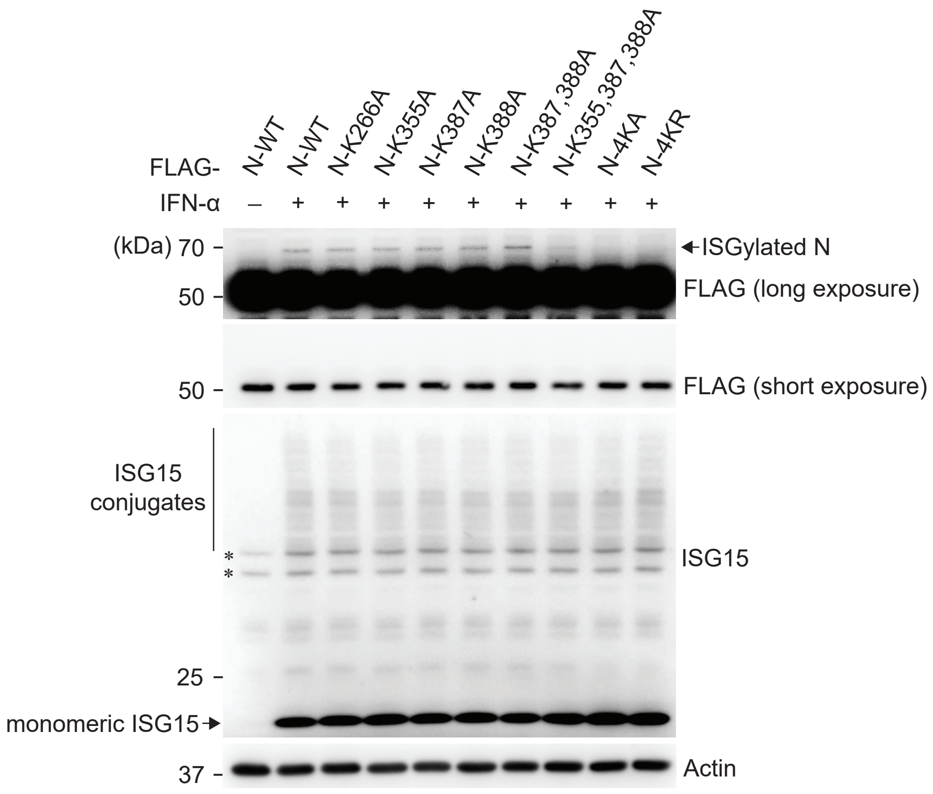


**Supplementary Figure 2**
